# Defining a Connectome-Based Predictive Model of Attentional Control in Aging

**DOI:** 10.1101/2021.02.02.429232

**Authors:** Stephanie Fountain-Zaragoza, Heena R. Manglani, Monica D. Rosenberg, Rebecca Andridge, Ruchika Shaurya Prakash

## Abstract

With advancing age, declines in the executive control of attention are accompanied by shifts in the functional topology of brain networks. However, there is increasing recognition of the considerable individual variability in the extent and types of attentional deficits that older adults exhibit, with results from neuroimaging investigations paralleling behavioral heterogeneity. Emerging computational methods leverage whole-brain functional connectivity to predict individual-level behaviors. These approaches are well-suited to the cognitive aging context, as they may elucidate configurations of functional connections that best explain group- and individual-level differences across older adults. Two independent samples of neurologically and psychiatrically healthy older adults were used to separately derive a predictive model of attentional control and test the model’s external validity. Here we show that despite challenges posed by heterogeneity in these aging samples, select functional connections carried meaningful variance, allowing for successful prediction of attention in a novel sample of older individuals.

## 1. Introduction

Age-related cognitive declines have far-reaching ramifications for older adults, a sector of the population that is rapidly growing from 55.7 million in 2016 to a projected 104.6 million by the year 2050 (U.S. Census Bureau, 2017). Even for those without clinical impairments, poorer cognitive function is associated with difficulties completing daily activities (Cahn-Weiner et al., 2000; Tucker-Drob, 2011), reduced quality of life (Netuveli et al., 2006; St. John & Montgomery, 2010), and increased mortality (Johnson, Lui, & Yaffe, 2007). These downward trends in cognition are thought to be explained by deficits in the executive control of attention, or the ability to select task-relevant information while inhibiting interfering information (Hasher & Zacks, 1988). Attentional control serves as a fundamental building block for many higher-order cognitive abilities that are implicated in daily functioning, thus representing a key area of interest in cognitive aging research. Importantly, within-person declines in attentional processes follow diverse trajectories over time (Goh et al., 2012, 2013), resulting in considerable individual variability in the magnitude and types of attentional deficits exhibited by older adults. Further, these deficits are not always evident in overt errors, and may instead manifest as compensatory strategy shifts observed only at the neural level (Lustig & Jantz, 2015). Thus, information represented in the brain’s functional architecture is of particular relevance to developing brain-based markers of attentional control in older adults.

Aging is known to have a prominent effect on the functional topology of the brain, characterized by reductions in the selectivity and specificity of neural recruitment. Older adults exhibit broad decreases in segregation between functional networks as well as disconnection within many networks (Liem et al., 2019). Changes in connectivity among canonical networks implicated in attentional control, such as the frontoparietal, salience, dorsal attention, and default mode networks, begin to appear as early as middle-age (Siman-Tov et al., 2017), and are observed across normal aging, mild cognitive impairment, and Alzheimer’s disease (Dennis & Thompson, 2014; Esposito et al., 2018; Sheline & Raichle, 2013). Age-related deficits in attentional performance have been linked to certain functional alterations, including decreased integrity of the default mode (Andrews-Hanna et al., 2007; Damoiseaux et al., 2008), frontoparietal (Geerligs, Renken, et al., 2015), and salience networks (Hausman et al., 2020; Onoda et al., 2012), as well as loss of segregation between the default mode network and the dorsal attention (Avelar-Pereira et al., 2017) and frontal executive control networks (Ng et al., 2016). However, the majority of previous studies have relied on select, *a priori* regions or canonical networks, likely missing critical variance represented across multiple neural systems.

In the last five years, there is growing recognition that complex cognitive constructs, such as attention, are an emergent property of functional interactions distributed across the whole brain. Thus, methods capitalizing upon connectome-wide patterns of connectivity may yield more comprehensive models of brain-behavior relationships that can be used to create informative predictive models (Woo et al., 2017). This study employed one such method, connectome-based predictive modeling (CPM; Shen et al., 2017), which uses cross-validation to identify functional networks that are predictive of behaviors of interest. CPM overcomes several limitations of previous approaches by employing a data-driven analysis that is not constrained to specific regions or networks and developing models that are informed by each individual’s unique connectivity features. This technique is thus particularly well-suited to the aging context, allowing us to move beyond brain-behavior associations to building models from brain features that predict cognition in previously unseen individuals.

Early application of CPM identified a model from task-based functional connectivity that captured significant variance in sustained attention performance in young adults using leave-one-out internal cross-validation (saCPM; Rosenberg, Finn, et al., 2016). This model generalized to several independent datasets, predicting attention-deficit/hyperactivity disorder symptoms in children from resting-state data (Rosenberg, Finn, et al., 2016) and predicting performance on multiple tasks of attention measuring inhibition and executive control from task-based data (Rosenberg et al., 2018; Rosenberg, Zhang, et al., 2016). These findings demonstrate that the CPM approach can yield markers of attentional ability that are generalizable across tasks and samples. In our work, we examined whether the saCPM model could further generalize to an aging sample, and demonstrated successful prediction of inhibitory control in a sample of both older and younger adults (Fountain-Zaragoza et al., 2019). However, connectivity within the saCPM did not account for significant age-related differences in performance, suggesting that it did not capture patterns of functional connectivity that are sensitive to the effects of aging.

The aim of the present study was thus to derive an age-specific connectome-based predictive model that explains individual differences in attentional control performance. Using a leave-one-out cross-validation technique, we attempt to identify patterns from whole-brain functional connectivity during a sustained attention task that predict sensitivity on that task in a sample of healthy older adults. We find that aging samples pose a unique challenge to predictive modeling as they exhibit considerable brain-behavior heterogeneity. Despite this, we identify an aging sustained attention CPM (Age-saCPM) and demonstrate its ability to predict attentional performance in an independent sample of older adults.

## 2. Material and Methods

### 2.1 Power Analysis

The primary aim of the study was to derive a functional connectivity-based marker predictive of attentional control performance in older adults. Network derivation was modeled after Rosenberg, Finn, et al., (2016), in which cross-validation was used to iteratively identify networks of connections (i.e., edges) that were related to attentional performance, generate predictive models, and test prediction on left-out participants. The resulting model (i.e., the saCPM) comprised a high-attention network, edges whose strength was associated with better performance (mean *r* = .59), and a low-attention network, edges whose strength was associated with worse performance (mean *r* = −.58). Based on an alpha level of .001 (two-tailed test), a total sample size of 40 participants was needed to yield an estimated power of at least 0.80 to derive networks for the CPM.

Additionally, our previous application of this model in a sample of older and young adults (Fountain-Zaragoza et al., 2019), found significant associations between saCPM predictions and observed Stroop task performance (*r*_s_= −.620, total sample size needed = 33). In order to have sufficient power for network identification, we derived networks using data from a set of 50 internal validation participants. We then assessed the external validity of the model on a previously unseen, independent sample of 34 participants.

### 2.2 Sampling and Screening

This cross-sectional study included data from two independent sets of healthy, community-dwelling older adults aged 65-85 years. Participants for the internal validation sample (*N*_1_ = 50) were recruited for a study aimed at identifying neuroimaging markers of attentional control in healthy older adults. This was the primary dataset used for network derivation, model building, and internal validation; participants were recruited for a study aimed at identifying biomarkers of healthy aging. The external validation sample (*N*_2_ = 34) was used to test prediction of the derived model; this sample was comprised of baseline data from an ongoing randomized controlled trial of mind-body interventions for healthy aging (clinical trial #NCT03626532). Participants were recruited from the greater Columbus, Ohio area.

Participants were considered eligible if they were right-handed, had corrected near and far visual acuity no worse than 20/40, were not colorblind (Ishihara, 2010), able to perceive and understand all study components, without objective cognitive impairment (see below), without contraindication to the MR environment, and self-reported absence of current psychiatric disorders, neurological disorders and incidents (e.g., stroke), and terminal illnesses (e.g., cancer). Participants were excluded if they scored in the range of possible mild cognitive impairment or dementia on cognitive screenings. This was defined as a score < 26 on the Montreal Cognitive Assessment (MoCA, Nasreddine et al., 2005; range: 26-30) for the internal validation sample. For the external validation sample, this was defined as a score < −1.5 *SD* on one or more of the following tests: the computerized Wisconsin Card Sorting Test (perseverations), WAIS-III Digit Span and Block Design subtests, Boston Naming Test, Controlled Oral Word Association Test, and the Hopkins Verbal Learning Test (total recall or retention). Both studies were approved by The Ohio State University Institutional Review Board, and informed consent was obtained from each participant. Neither dataset has been published. Data supporting the findings of this study can be made available upon request from the corresponding author.

### 2.3 Data Exclusion

Of the 50 internal validation participants, 41 had usable data. Of the 34 external validation participants, 26 had usable data. Reasons for data exclusion included acquisition error (*N*_1_: 1), possible mild cognitive impairment (*N*_2_: 1), incidental findings on structural images (*N*_1_: 2; *N*_2_: 1), poor engagement with the gradual-onset continuous performance task (gradCPT hit rate >2.5 SD below the mean, translating to 52%-67% correct; *N*_1_: 2), inadequate vision correction (*N*_1_: 1), low correlation between their functional connectivity matrix and all other functional connectivity matrices in the sample (−2.7 SD below the mean; *N*_1_: 1), excessive head motion throughout both runs of the gradCPT (*N*_1_: 2; *N*_2_: 6). Of the 41 internal validation participants, six demonstrated excessive motion in the second run of the gradCPT, so only the first run was included in the analyses. Of the 26 external validation participants, three had only one run included in analyses due to excessive motion, misinterpreting task instructions, and a low hit rate. Consistent with a well-established pipeline in our laboratory (Fountain-Zaragoza et al., 2019) and a prior study employing CPM in older adults (Lin et al., 2018), excessive head motion was defined as mean framewise displacement (FD) > .15 mm and motion > .5 mm in more than 10% of functional volumes (Power, Barnes, Snyder, Schlaggar, & Petersen, 2012).

The distribution of each variable of interest (*d’*, connectivity strength for each network, predicted performance generated from each network, and mean FD) was checked using the Shapiro-Wilk test of normality (Shapiro & Wilk, 1965). Of these variables, mean FD was not normally distributed.

### 2.4 Image Acquisition and Analysis

#### 2.4.1 Gradual-onset Continuous Performance Task

While in the scanner, participants performed the gradual-onset continuous performance task (gradCPT; Esterman et al., 2013; Fortenbaugh et al., 2018; Rosenberg, Finn, et al., 2016). In this task, participants viewed grayscale, circular images (diameter = 256 pixels, 7° of visual angle) of city and mountain scenes presented in the center of the screen. Images were presented one at a time. Images gradually transitioned using linear pixel-by-pixel interpolation, taking 800 ms for each full transition. Participants were asked to respond by pressing with their right index finger to city scenes, occurring randomly about 90% of the time, and to withhold responses to mountain scenes. They completed two separate runs, each consisting of four 3-minute blocks interleaved with rest blocks. Rest blocks were indicated by the presentation of a circle in the center of the screen for 30 seconds, during which participants were asked to simply attend to the fixation circle. They were then alerted to the start of the next block by the presentation of a dot inside the circle for an additional 2 seconds. Eight seconds of fixation were included at the start of each run and were excluded from analyses. Each run lasted 13:44 minutes. The dependent variable of interest was sensitivity, or *d’*, which takes both correct responses (i.e., hits) and errors of commission (i.e., false alarms) into account. This was calculated as *d’ = z*(hit rate)–*z*(false alarm rate) for each block and then averaged across blocks for each participant (as in Rosenberg, Finn, et al., 2016). Reliability of *d’* was calculated using a Spearman-Brown-corrected split-half correlation between average performance on even blocks versus odd blocks. There was excellent reliability in both the internal and external validation samples (*r*s = 0.97).

#### 2.4.2 Imaging Parameters

Data for both studies were collected at the Center for Cognitive and Behavioral Brain Imaging at The Ohio State University on a 3T Siemens MAGNETOM Prisma system using a 32-channel head coil. The order of sequence acquisition for the internal validation sample (N_1_) was: localizer, resting-state, T2, gradCPT run 1, magnetization prepared rapid gradient echo (MPRAGE), fieldmaps, gradCPT run 2. The order of sequence acquisition for the external validation sample (N_2_) was: localizer, resting-state, N-back task run 1, N-back task run 2, MPRAGE, gradCPT run 1, T2, fieldmaps, gradCPT run 2, DTI. This study required the use of at least one of the gradCPT runs, the MPRAGE for registration of functional images, and the fieldmaps for distortion correction. Sequence parameters for the gradCPT were identical for both samples. For each run of the gradCPT task, 824 volumes were acquired using a multiband echo-planar imaging (EPI) sequence with the following parameters: repetition time (TR) = 1000 ms, echo time (TE) = 28 ms, flip angle = 50°, field of view: 240 mm x 240 mm, 45 axial slices, slice thickness = 3 mm (voxel size = 3.0 mm^3^), multiband acceleration factor = 3. For the internal validation sample (N_1_), the anatomical MPRAGE had the following parameters: TR = 1900 ms, TE = 4.44 ms, flip angle = 12°, field of view: 256 mm x 256 mm, slice thickness = 1.00 mm (voxel size = 1.0 mm^3^), 176 sagittal slices. For the external validation sample (N_2_), MPRAGE parameters were: TR = 2400 ms, TE = 2.15 ms, flip angle = 8°, field of view: 256 mm x 256 mm, slice thickness = 1.00 mm (voxel size = 1.0 mm^3^), 208 sagittal slices. For both samples, the fieldmap had the following parameters: TR = 512 ms, TE 1 = 5.19 ms, TE 2: 7.65 ms, flip angle = 60°, field of view: 240 mm x 240 mm, slice thickness = 3.00 mm (voxel size = 2.0 × 2.0 × 3.0 mm), 45 axial slices.

#### 2.4.3 Image Preprocessing

Data were preprocessed using fMRIB’s software library (FSL; Smith et al., 2004) in two steps. First, a minimal preprocessing pipeline was used that included motion correction, brain extraction, bias field inhomogeneity correction, and spatial smoothing with a 6-mm kernel. Second, all steps requiring linear regression with projection of data into an orthogonal space were conducted in a joint nuisance regression model (as recommended in Lindquist, Geuter, Wager, & Caffo, 2019). This nuisance regression step included pre-whitening and high-pass filtering (frequency = 0.01 Hz) to remove the effects of low-frequency noise, and regression of mean signal from CSF and WM, a 24-parameter motion model (six motion parameters, six temporal derivatives, and their squares), and mean global signal, which has been found to reduce the effects of both physiological signals and head motion in functional connectivity data (Ciric et al., 2017; Parkes et al., 2018; Power et al., 2014; Power, Plitt, Laumann, & Martin, 2017; Power, Schlaggar, & Petersen, 2015; Weissenbacher et al., 2009). Additionally, following the recommendation of Power et al. (2012), volumes with FD values > 0.5 mm were added as confounds in the regression model. See supplemental information for more details regarding motion controls.

#### 2.4.4 Whole-Brain Functional Connectivity

Brain parcellation was conducted using a 268-node, whole-brain functional atlas (Shen, Tokoglu, Papademetris, & Constable, 2013; https://www.nitrc.org/frs/?group_id=51). This atlas includes cortical, subcortical, and cerebellar regions and was developed with the goal of maximizing the similarity of voxel-wise timeseries within each node. Nodes were considered missing if less than 50% of their original volume was retained after masking each participant’s functional image, resulting in only two excluded nodes (nodes #51: right temporal pole, node #252: left cerebellum from the Shen atlas). Whole-brain functional connectivity during task runs was computed for each participant using the Graph Theory GLM toolbox (GTG; Spielberg, Miller, Heller, & Banich, 2015). The first eight volumes of rest from each run, as well as embedded rest breaks, were excluded and remaining data were concatenated across runs for participants with two usable runs. Activity in each node was calculated by averaging the time courses of all voxels within the node. Functional connectivity (i.e., edges) was calculated as the Pearson correlation (*r*) between the timecourses of each node pair. Each coefficient was then normalized using Fisher’s *r*-to-*z* transformation and the resulting 266 × 266 fully connected matrices representing the functional connectivity between each node pair were used in all subsequent analyses.

#### 2.4.5 Effects of Head Motion

Given that motion can introduce significant confounds in studies of functional connectivity, after excluding participants with excessive head motion and correcting for head motion during preprocessing, we assessed whether estimates of head motion (mean FD), derived from processed data, were correlated with *d’*, age, or the strength of the high- or low-attention networks following derivation. Head motion was significantly associated with lower *d’* (*r*_s_ = −0.31, *p* = .048), but not age (*r*_s_ = 0.25, *p* = .12), in the internal validation sample. As such, mean FD was used as a covariate at the edge selection step in internal validation. In the external validation sample, motion was not significantly associated with *d’* (*r*_s_ = 0.25, *p* = .22) or age (*r*_s_ = −0.096, *p* = .64) so neither were included as covariates in the primary analyses. However, to be conservative, the external validation effects were additionally tested when controlling for head motion.

### 2.5 Statistical Analyses

#### 2.5.1 Predicting Attentional Control

In all subsequent analyses assessing the performance of predictive models, we report several metrics of model performance. These include Spearman rank correlation between observed and predicted *d’*, mean squared error (MSE) from the linear regression fitting predicted to observed *d’*, and the coefficient of determination prediction *R*^2^ (also known as *q*^2^; Scheinost et al., 2019). Prediction *R*^2^ is computed as [1-(model MSE/null MSE)] in which a null model has predictions that are all the same (the mean of observed behavior). Due to this, prediction *R*^2^ can be negative, indicating that the predictive model explained less variance than simply predicting the mean of *d’*.

#### 2.5.2 Predictive Ability of the saCPM in Aging Datasets

We first evaluated the predictive ability of the young-adult saCPM (Rosenberg, Finn, et al., 2016) in both samples of older adults separately (*N*_1_ = 41 and *N*_2_ = 26). To do so, we applied the masks from the original saCPM high-attention and low-attention networks (268 × 268 symmetrical, binary matrices including 1s for edges that belong in the network and 0s elsewhere) to individual functional connectivity matrices. We averaged all edge values in each mask to calculate high-attention and low-attention connectivity strengths and then computed combined strength (i.e., the different between high- and low-attention network strength) for each individual. Network strength values were entered into the GLM model that was constructed in the initial derivation of the saCPM (Rosenberg, Finn, et al., 2016) to generate predicted *d’* for all participants. Model performance was assessed as the correspondence between observed and predicted *d’* using the metrics specified above.

#### 2.5.3 Internal Validation

The CPM method, previously described in (Finn et al., 2015; Rosenberg, Finn, et al., 2016; Shen et al., 2017), was used to predict individual differences in attentional control (i.e., *d’* on the gradCPT) from whole-brain functional connectivity. All CPM-based analyses were completed using custom MATLAB scripts (adapted from those available here: https://github.com/YaleMRRC/CPM). We defined the models using a leave-one-out cross-validation (LOOCV) approach to reduce the rates of false-positive findings. On each iteration of cross-validation, the model was trained on data from *n*-1 participants (*n* = 40) and tested on the left-out individual. Forty-one rounds of cross-validation were conducted such that each individual served as the test participant once. Each iteration included feature selection followed by model building and prediction of performance for the left-out individual.

In the feature selection step, we identified edges that were related to *d’*. Partial Pearson correlations were conducted between every edge and *d’* across training set participants, covarying mean FD. These correlation coefficients were then thresholded at *p* < .01 across participants and separated into a positive tail called the high-attention network (i.e., edges whose strengths are associated with better performance) and a negative tail called the low-attention network (i.e., edges whose strengths are associated with poorer performance). We then computed a single summary measure (strength) for each network by averaging all constituent non-zero edges, which represents each participant’s level of functional connectivity within the respective set of edges. We consolidated the information from the high- and low-attention networks using combined network strength, calculated as the difference between network strengths. In order to facilitate interpretability, results are presented for this combined network model, unless otherwise stated.

We next built three predictive models based on network strengths within the Aging high-attention network, Aging low-attention network, and the combined model. For each, we used linear regression to generate a first-degree polynomial that best fit respective network strength to *d’* in the set of *n* – 1 (i.e., 40) participants. We then tested prediction on the left-out participant by calculating their three network strengths and entering those values into each model to generate three predicted *d’* scores. This process was repeated for all iterations of *n* – 1 groups, such that predicted scores were eventually generated for each participant. The final high- and low-attention networks of this new Age-saCPM is the set of edges that appeared in every round of LOOCV.

Model success was quantified by conducting a Spearman rank correlation (*r*_s_) between predicted and observed *d’*. Model fit indices, such as MSE and prediction *R*^2^, were not reported for internal validation results because in-sample predictive accuracy likely overestimates the effect (Poldrack, Huckins, & Varoquaux, 2019). We chose to use Spearman rank correlations because CPM predictions have been shown to have limited range compared to observed scores, and are thus better represented as relative rather than absolute predictions (Rosenberg, Finn, et al., 2016). Given the lack of independence across LOOCV iterations, permutation testing was used to determine significance of any successful models based on a non-parametric *p*-value calculated from 1,000 permutation tests. To do so, we randomly shuffled participants’ *d’* values and conducted LOOCV to assess the correlation between predicted and randomly shuffled observed scores. This was done 1,000 times in order to generate a null distribution of correlation values, against which the results of the analysis were compared to determine significance level (i.e., *p* value of each network).

#### 2.5.4 External Validation

We next assessed the predictive power of the model by correlating predicted and observed *d’* for a previously unseen external validation set. Network strength values for each of the external validation participants were computed by averaging all edge values within the final high- and low-attention network masks generated in the previous step. We also computed combined network strength by subtracting the low-attention network strength from the high-attention network strength. Network strength values were entered into the GLM models to produce predicted *d*’ values. Model performance was assessed as the correspondence between observed and predicted *d’* using the metrics specified above.

#### 2.5.6 Characterizing the Anatomy of the Age-saCPM

We characterized the distribution of edges within the Age-saCPM using two approaches to classify edges. First, we used matrices to visualize how many edges in the high-attention and low-attention networks belonged to each macroscale brain region including prefrontal, motor strip, insula, parietal, temporal, occipital, limbic, cerebellum, subcortical, and brainstem. Next, we calculated the relative involvement of ten canonical networks in the Age-saCPM. The networks were previously defined in Finn et al. (2015) and Noble et al. (2017) and include: 1) medial frontal (MF), 2) frontoparietal (FP), 3) default mode (DMN), 4) motor (MOT), 5) visual I (V1), 6) visual II (V2), 7) visual association (VA), 8) salience (SAL), 9) subcortical (SC), and 10) cerebellar (CB). We employed a calculation to index the relative contribution of each network, as reported by Greene et al. (2018). We designated each edge (*i*,*j*) to its respective pair of canonical networks [A (the network that includes *i*), B (the network that includes *j*)]. Conceptually, the contribution of edges between canonical networks A and B is the number of edges between A and B belonging to the Age-saCPM (*m*_A,B_) divided by the number of edges between A and B in the whole brain (*E*_A,B_). These two numbers were also normalized for network size (*m*_tot_ = total number of edges in the Age-saCPM; *E*_tot_ = total number of edges in the whole brain), such that a value of 1 indicates a proportionate contribution and larger values indicate a greater contribution of that network pair to the model than would be expected for its size. The resulting matrices are symmetric along the diagonal, so only the bottom triangle of each matrix is displayed.

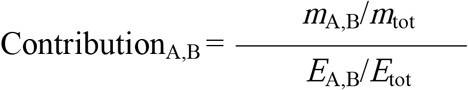

## 3. Results

### 3.1 Predicting Attentional Control

Two independent groups of older adults (internal validation sample: *N*_1_ = 41, external validation sample: *N*_2_ = 26) performed the gradual onset continuous performance task (gradCPT; Esterman et al., 2013; Fortenbaugh et al., 2018; Rosenberg, Finn, et al., 2016) in the scanner, and performance was measured using sensitivity (*d’*). Task-based functional connectivity was calculated using the 268-node Shen functional atlas (Shen et al., 2013; https://www.nitrc.org/frs/?group_id=51). The two included samples did not significantly differ on demographic or beahvioral variables (Table 1).

**Table 1.** Demographics and Descriptive Statistics.

| | Internal Validation (full $n = 50$) | | External Validation (full $n = 34$) | | Internal Validation (included $n = 41$) | | External Validation (included $n = 26$) | | Group Difference (included) | |
| --- | --- | --- | --- | --- | --- | --- | --- | --- | --- | --- |
| | Mean | SD | Mean | SD | Mean | SD | Mean | SD | test | $p$ |
| **Demographics** |  |  |  |  |  |  |  |  |  |  |
| Age (years) | 70.3 | 4.60 | 70.7 | 4.53 | 69.7 | 3.88 | 69.5 | 3.41 | $t(65) = 0.16$ | .88 |
| Education (years) | 17.4 | 3.21 | 16.5 | 2.77 | 17.5 | 3.30 | 16.2 | 2.32 | $t(65) = 1.73$ | .08 |
| % Women | 54.0 | --- | 47.1 | --- | 51.2 | --- | 46.2 | --- | $X^2 = 0.16$ | .69 |
| Race (no. and %) | | | | | | | | | $X^2 = 1.39$ | .71 |
| White | 47 | 94.0 | 31 | 91.2 | 38 | 92.7 | 25 | 96.2 |  |  |
| Black | 1 | 2.00 | 3 | 8.82 | 1 | 2.44 | 1 | 3.85 |  |  |
| Asian | 1 | 2.00 | 0 | 0.00 | 1 | 2.44 | 0 | 0.00 |  |  |
| Other | 1 | 2.00 | 0 | 0.00 | 1 | 2.44 | 0 | 0.00 |  |  |
| **MRI Assessment** |  |  |  |  |  |  |  |  |  |  |
| gradCPT $d'$ | 2.59 | 0.75 | 2.22 | 0.87 | 2.72 | 0.70 | 2.47 | 0.62 | $t(65) = 1.46$ | .15 |
| Raw meanFD | 0.17 | 0.057 | 0.23 | 0.12 | 0.17 | 0.048 | 0.18 | 0.04 | $t(65) = -1.47$ | .15 |
| Included volumes (no. and %) | 1598.6 | 97.0 | 1509.2 | 91.6 | 1610.3 | 97.7 | 1596.6 | 96.9 | $t(65) = 0.98$ | .33 |

#### 3.1.1 Predictive Ability of the saCPM in Aging Datasets

In our previous work, we found that the original saCPM, identified in a sample of healthy young adults (Rosenberg, Finn, et al., 2016), predicted inhibitory control (Stroop task performance) across a sample of both older and younger adults (Fountain-Zaragoza et al., 2019). However, we found that prediction within older adults alone was only successful from the low-attention network, and only a small subset of edges within it (the OA Low1 and OA Low2 subnetworks) accounted for age-related performance differences. Given this mixed evidence of generalizability to aging samples, we tested the predictive ability of the original saCPM model and age-sensitive subnetworks in the two older-adult samples used in this study. This allowed a more direct evaluation of generalizability, as these participants performed the same attention task on which the original saCPM was derived (i.e., the gradCPT).

When the saCPM was applied to the internal validation dataset (*N*_1_ = 41), prediction was not successful from the high-attention network (*r*_s_ = 0.024, *p* = .88, MSE = 0.51, prediction *R*^2^ = −4.86%), the low-attention network (*r*_s_ = −0.13, *p* = .43, MSE = 0.50, prediction *R*^2^ = −2.87%), or the combined model (*r*_s_ = −0.057, *p* = .72, MSE = 0.50, prediction *R*^2^ = −4.47%). Applying the saCPM to the external validation dataset (*N*_2_ = 26), predictions were also not successful from the high-attention network (*r*_s_ = −0.21, *p* = .29, MSE = 0.38, prediction *R*^2^ = −1.50%), the low-attention network (*r*_s_ = −0.32, *p* = .11, MSE = 0.38, prediction *R*^2^ = −2.33%), or the combined model (*r*_s_ = −0.34, *p* = .087, MSE = 0.37, prediction *R*^2^ = 0.077%). The negative prediction *R*^2^ values indicate that saCPM predictions were less accurate than simply predicting the mean of observed *d’* in each sample. Similarly, connectivity strength wtihin the age-sensitive saCPM subnetworks was not significantly associated with *d’* in either the internal validation sample (OA Low1 subnetwork: *r*_s_ = −0.091, *p* = .57; OA Low2 subnetwork: *r*_s_ = −0.024, *p* = .88) or the external validation sample (OA Low1 subnetwork: *r*_s_ = 0.19, *p* = .34; OA Low2 subnetwork: *r*_s_ = 0.061, *p* = .77). The lack of successful prediction from existing networks to both older-adult samples suggests that there are limits to the generalizability of the saCPM networks in aging samples and further justifies the need for defining an aging-specific model of attentional control.

#### 3.1.2 Internal Validation

Connectome-based predicitve modeling (CPM; Shen et al., 2017) was conducted using the internal validation sample (N_1_) with the goal of identifying a set of functional connections that predicts performance on the gradCPT (i.e., *d’*). Network models were iteratively trained using leave-one-out cross-validation (LOOCV) on 40 participants’ task data and tested on the left-out 41^st^ participant. On each round, networks were derived by identifying edges (i.e., functional connections) that were significantly associated (at *p* < .01) with *d’*, controlling for head motion, in the 40 training set participants. Strength in the identified network was summarized as mean connectivity among included edges, and a linear model was used to relate network strength to behavior across the training set. Then, the left-out participant’s network strength value was entered into the linear model to generate a predicted *d’* score. Model performance was assessed by correlating individuals’ observed *d’* with their predicted *d’* generated during the respective round of LOOCV. Network derivation did not result in significant prediction of left-out participants’ *d’* from the high-attention network (*r*_s_ = 0.07, *p* = .66), the low-attention network (*r*_s_ *o* = 0.25, *p* = .11), or a combined model (*r*_s_ = 0.16, *p* = .31), calculated by subtracting low-attention network strength from high-attention network strength. Given that internal validation was unsuccessful, permutation testing was not conducted.

We tested the influence of several confounding variables on derivation, including residual effects of head motion, cross-validation and modeling strategy, and choice of significance threshold. We confirmed that the models were not fitting to noise attributable to motion (see Supplementary Materials “Additional Motion Controls”). Derivations remained non-significant at multiple thresholds based on both *p*-values and percentiles (see Supplementary Materials “Testing Edge Selection Thresholds” and Table S1). We also found that prediction remained unsuccessful across 100 permuted iterations of 10-fold cross-validation and when using LOOCV with ridge regression (see Supplementary Materials “Alternative Derivation Strategies and Figure S1).

#### 3.1.3 Examining Cross-Round Consistency

In order to investigate why internal validation may have failed, we assessed the consistency of edge selection across rounds of LOOCV by correlating the two-dimensional matrix of edges from one round with the matrices of each of the other rounds. Across rounds, the correlation strengths ranged from 0.55 −0.86 (*M* = 0.79, *SD* = 0.066) and the average percent overlap ranged from 62% to 87% (*M* = 79%, *SD* = 6%). These two metrics suggest that there was considerable variability in the edges selected across rounds of LOOCV (i.e., a small number of common edges), and raised the possibility that the model was overfitting to noise. Therefore, as predictive reliability is contingent upon selecting correlated features that capture variance both in the construct of interest and between individuals (Sripada et al., 2020), unsuccessful internal predictions may have been driven by inconsistent or non-specific feature selection. Additionally, the range of predicted *d’* in this study was smaller (range = 1.22, min = 2.23, max = 3.45) than the range of observed scores (range = 2.53, min = 1.54, max = 4.07), such that the model did not generate predicted scores below −0.69 *SD* or above 1.04 *SD* from the observed *d’* mean. As such, prediction error was smallest (i.e., the model was most accurate) for individuals who scored within 1 *SD* of the mean. Although this phenomenon is observed in both CPM-specific analyses (e.g., Rosenberg, Finn, et al., 2016), and machine learning predictions in general, these observations suggest that patterns of functional connectivity-performance associations may differ for individuals performing on the tails of the distribution compared to the rest of the sample. Thus, failure to accurately predict performance across rounds of LOOCV may also be due to heterogeneity in brain-behavior relations.

#### 3.1.4 External Validation

Establishing the validity of predictive models requires assessment of model performance in an independently collected dataset that was not involved in model building (Scheinost et al., 2019). This process ensures that model predictions are valid and generalizable, rather than overfitting to sample-specific features (Yarkoni & Westfall, 2017). In this case, internal validation was hampered by variability in edges selected across rounds of cross-validation. However, construction of a final model, containing edges appearing in every round of cross-validation, mitigates this problem and can be used to assess the external validity of the Age-saCPM in a completely independent external validation dataset (*N*_2_). Final networks were defined as edges found consistently across rounds of internal cross-validation and GLM coefficients were calculated by modeling the relationship between connectivity in these final networks and behavior across the full internal validation sample. The final Age-saCPM contained 122 edges; 71 in the high-attention network and 51 in the low-attention network (Figure 1a).

**Figure 1.**
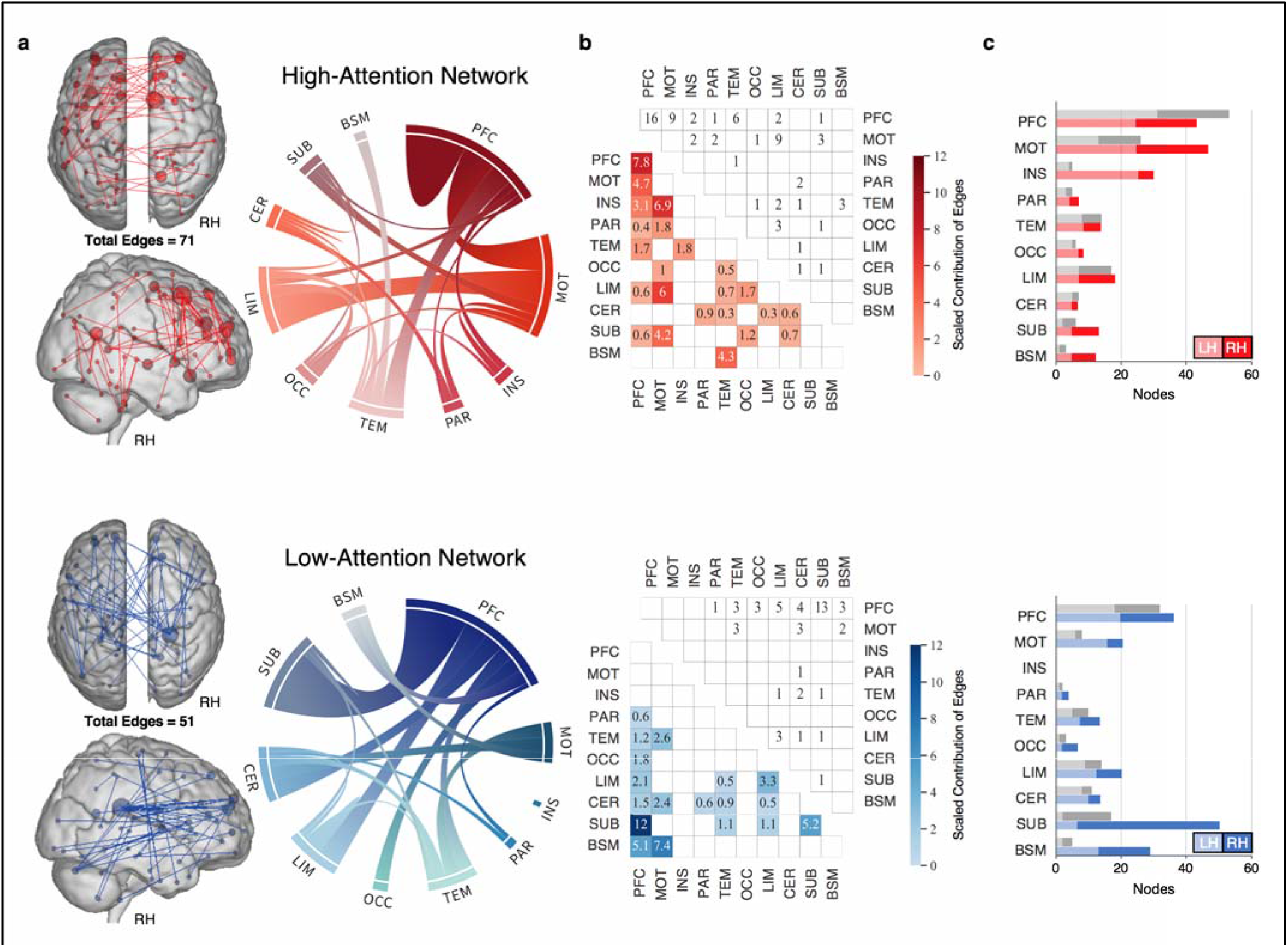
Anatomical Distribution of the Age-saCPM. A) The 71 edges belonging to the high-attention network are in red and the 51 edges belonging to the low-attention network are in blue. In brain figures, the size of nodes represents higher degree (i.e., more functional connections). Ring plots depict intra- and inter-region connections for the high-attention (red) and low-attention (blue) networks. Ribbon size is proportional to the number of edges. For B and C, scaled contributions were calculated by adjusting for the number of edges belonging to the respective regions and the size of the networks. Values > 1 indicate a disproportionate contribution relative to size. B) Matrices present the contributions (lower triangular) and raw number of edges (upper triangular) for each macroscale brain region for the high-attention (red) and low-attention (blue) networks. C) Bars represent the nodes in the high-attention (red, top) and low-attention (blue, bottom) networks belonging to each macroscale brain region in the left hemisphere (lighter shades) and right hemisphere (darker shades). Colored bars depict contributions (values multiplied by 10 for visualization purposes), gray bars represent the raw number of nodes. Region acronyms: PFC = prefrontal; MOT = motor; INS = insula; PAR = parietal; TEM = temporal; OCC = occipital; LIM = limbic; CER = cerebellum; SUB = subcortex; BSM = brainstem. Visualization software: glass brains (https://bioimagesuiteweb.github.io/webapp/connviewer.html), ring plots (https://flourish.studio/).

As a true test of generalizability, we examined the predictive power of this final Age-saCPM model in the independent external validation sample of older adults who also completed the gradCPT (Figure 2). We found that prediction was successful from the combined Age-saCPM networks (*r*_s_ = 0.55, *p* = .004, MSE = 0.30), accounting for 18.7% (prediction *R*^2^) of the variance in *d’* in previously unseen individuals, and this effect remained when controlling for head motion (*r*_s_ = 0.51, *p* = .009). The low attention network accounted for more variance (*r*_s_ = 0.60, *p* = .002, MSE = 0.28, prediction *R*^2^ = 24.8%) than the high-attention network (*r*_s_ = 0.39, *p* = .052, MSE = 0.35, prediction *R*^2^ = 6.55%). This pattern is consistent with our previous finding that the original saCPM networks identified by Rosenberg, Finn, et al. (2016) generalized to an older adult sample and that components of the low-attention network were more sensitive to age-related deficits in inhibitory control (Fountain-Zaragoza et al., 2019). Notably, prediction in the external validation dataset was successful from models derived using all thresholds tested (Supplementary Figure S2). However, the *p* < .01 model demonstrated greater predictive power than models with more lenient thresholds, and stricter models yielded very low numbers of edges (ranging from 9-58).

**Figure 2.**
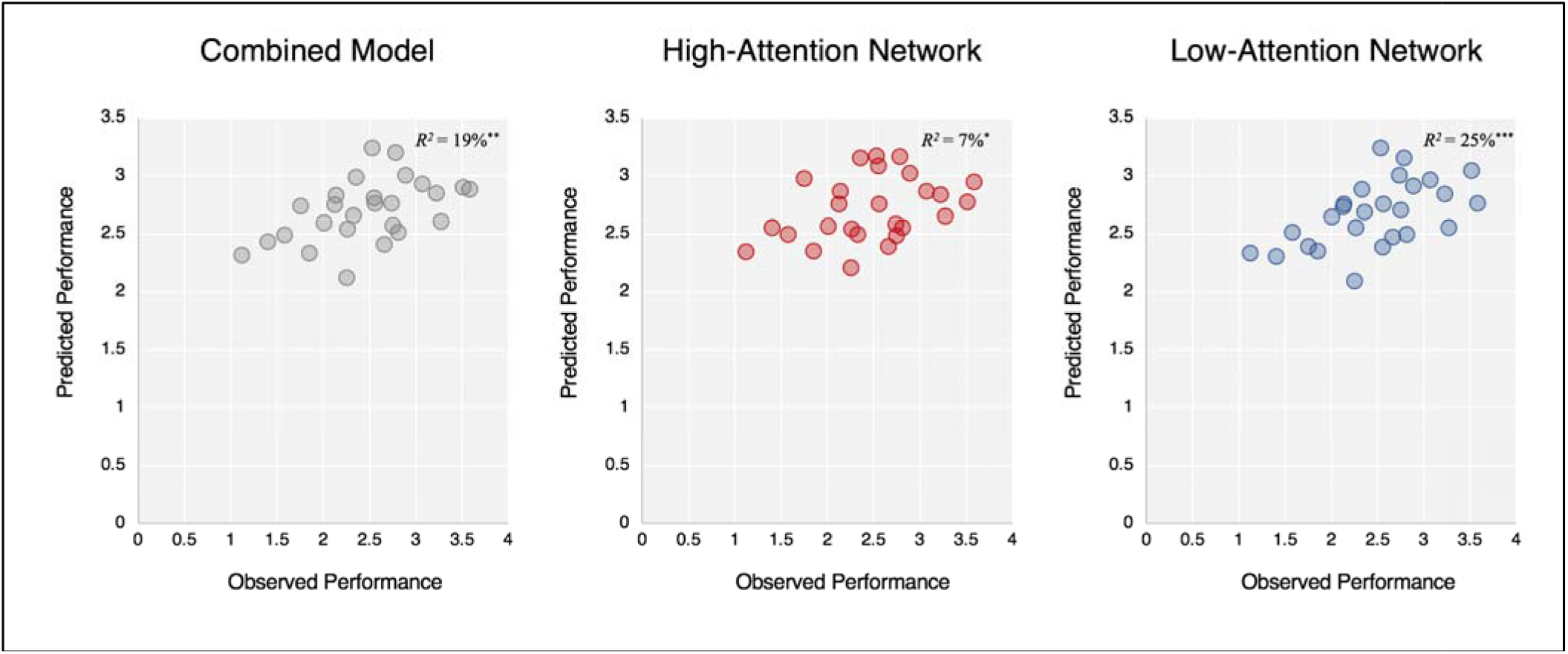
The Age-saCPM Predicts Attentional Performance in Novel Older Adults. Scatter plots show the correspondence between observed performance on the gradCPT (*d’*) and model-based predicted performance based on the combined network (gray), high-attention network (red), and low-attention network (blue). The coefficient of determination (prediction *R*^2^) is presented for each. \**p* < .05, \*\**p* < .01, \*\*\**p* < .001.

#### 3.1.5 Characterizing the Anatomy of the Age-saCPM

The distribution of Age-saCPM edges and nodes belonging to each macroscale region of the brain are presented in Figure 1a-c. Of the 71 edges in the high-attention network, intra-prefrontal edges represented the largest proportion (22.5%) and had the largest contribution to the network, accounting for size. Others with large contributions included insular-motor, limbic-motor, and prefrontal-motor edges. Contrastingly, over a quarter of the 51 edges in the low-attention network were between prefrontal and subcortical areas (25.5%), representing the largest contribution to the network. This was followed by contributions from motor-brainstem edges, prefrontal-brainstem edges, and intra-subcortical edges (Figure 1b). As can be seen in Figure 1c, the most striking difference between the two networks was the much greater involvement of bilateral prefrontal nodes in the high- than low-attention networks. The observed bilateral, and particularly left hemisphere, involvement fits with a pattern of age-related neural compensation in response to demanding tasks, which has been well-established in regional activation studies (Grady, 1998; Park & Reuter-Lorenz, 2009; Reuter-Lorenz & Cappell, 2008). This is also consistent with previous functional connectivity studies that have demonstrated compensation such that older adults who demonstrated increased bilateral prefrontal connectivity performed similarly to young-adult comparators, whereas the group that did not performed much lower (Eavani et al., 2016). The low-attention network had greater representation of subcortical and cerebellar nodes, which are increasingly recognized as playing important roles in both motor and attentional functioning (Buckner, 2013; Kellermann et al., 2012; Strick, Dum, & Fiez, 2009). In fact, certain cerebellar regions participate as nodes in the dorsal attention network (Brissenden, Levin, Osher, Halko, & Somers, 2016). However, aging is associated with reduced cerebellar volume and altered cortico-cerebellar network connectivity, both of which have been linked to age-related performance decrements (Bernard & Seidler, 2014). Thus, the involvement of subcortical and cerebellar regions in predicting poorer performance may reflect age-related neural vulnarbilities in those areas.

Of note, there was very limited overlap between the Age-saCPM and the original saCPM networks (Rosenberg, Finn, et al., 2016) with just one edge overlapping in the high-attention networks, zero in the low-attention networks, and 4 edges when network membership is ignored. There was no overlap between the Age-saCPM and saCPM subnetworks reported in Fountain-Zaragoza et al. (2019).

The relative involvement of each of ten canonical functional networks in the Age-saCPM is depicted in Figure 3. The greatest contributions to the high-attention network were from connections among the medial frontal and frontoparietal networks and within the salience network. In addition to being consistent with the patterns of compensatory frontal recruitment discussed above, this pattern highlights the non-specific nature of this recruitment such that we observed connectivity between multiple frontal networks. This is consistent with findings of age-related reductions in network segregation, particularly among networks implicated in associative processes, including frontoparietal, salience, and dorsal/ventral attention networks (Chan et al., 2014). Our results suggest that interactions among these frontal networks supports attentional performance for older adults. This pattern also mirrors results from previous functional connectivity-based predictive modeling studies that highlight an important role of these networks for individual-level performance. One paper found that medial frontal and frontoparietal network connectivity was most successful at discriminating between individuals (Finn et al., 2015) and another showed that predictive features for a variety of cognitive tasks (including working memory, language, and motor tasks) were concentrated in medial frontal, frontoparietal, visual, and motor networks (Greene et al., 2020). For the low attention network, there was greatest involvement of connections between the visual II and medial frontal networks and between subcortical and medial frontal networks. Although somewhat unexpected, it is possible that the involvement of medial frontal-visual II connectivity in predicting poorer performance reflects failed attempts to engage with the visual stimuli and cognitive demands of the task. The involvement of subcortical-frontal connectivity, on the other hand, is consistent with results observed in the original saCPM, in which there were more connections within the subcortical-cerebellum network and between subcortical-cerebellum and frontoparietal networks in the low-attention than high-attention networks (Rosenberg, Finn, et al., 2016). This pattern may reflect age-related increases in connectivity from subcortical regions to other networks (Tomasi & Volkow, 2012), and might suggest a detrimental role of these shifts for attentional performance.

**Figure 3.**
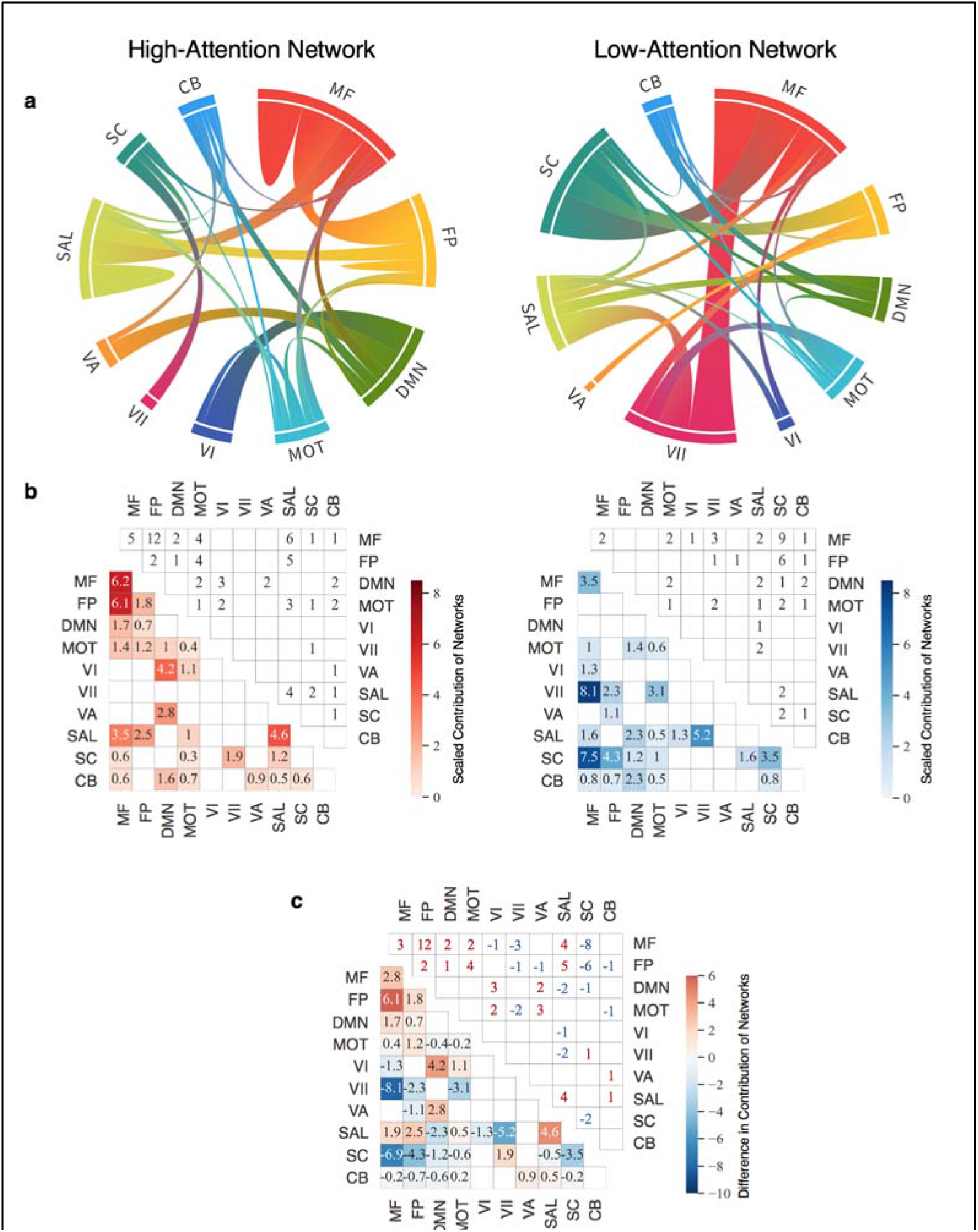
Functional Distribution of the Age-saCPM. A) The involvement of ten canonical networks in the high-attention (left) and low-attention (right) networks. Ribbon size is proportional to the degree of contribution of each network pair, which adjusts for the number of edges belonging to the respective networks and the size of the Age-saCPM networks. Values > 1 indicate a disproportionate contribution relative to size. B) Matrices present the contributions (lower triangular) and raw number of edges (upper triangular) for each canonical network pair in the high-attention (red) and low-attention (blue) networks. C) The difference in contribution (lower triangular) and number of edges (upper triangular) of each pair of canonical networks. Red represents higher involvement in High > Low, blue represents higher involvement in Low > High. Network acronyms: MF = medial frontal; FP = frontoparietal; DMN = default mode; MOT = motor; VI = visual I; VII = visual II; VA = visual association; SAL = salience, SC = subcortical; CB = cerebellar. Ring plot visualization: https://flourish.studio/.

## 4. Discussion

This study aimed to define a functional connectivity-based whole-brain model of attentional control in healthy older adults. A network-based predictive model previously defined in young adults (the saCPM; Rosenberg, Finn, et al., 2016) did not generalize to the two samples of older adults in this study, demonstrating the need for an aging-specific model. Using connectome-based predictive modeling, our initial attempt to identify brain networks that were predictive of attentional performance was unsuccessful. We observed that the consistency of edges selected across rounds of cross-validation was low, which appeared to be driven by heterogeneity in brain-behavior relationships across participants. However, the final network of edges that appeared in every round of cross-validation successfully predicted performance in an independent sample of older adults. Thus, despite the observed heterogeneity, there is a shared network of connections that predicts meaningful variance relevant to attention in older adults.

The final Age-saCPM model successfully predicted attentional control from task-based functional connectivity in an independent sample of healthy older adults, accounting for roughly 25% of the variance in performance. Prediction in this external sample was stronger from the low-attention network (24.8% of the variance) than the high-attention network (6.5% of the variance). This pattern is consistent with our previous finding that components of the low-attention network from the original saCPM were more sensitive to age-related deficits in inhibitory control (Fountain-Zaragoza et al., 2019). Interestingly, the original young-adult saCPM networks (Rosenberg, Finn, et al., 2016) did not successfully predict performance in either sample in this study, suggesting that a model trained in older adults may contain signal that is specific to aging and thus offer superior generalizability in this population. Further, we found little overlap between the original saCPM and the Age-CPM, with only four shared edges between the two. The distinctiveness of the Age-CPM from previous models derived in young adults may be driven by a unique attentional signature in aging.

The networks of the Age-saCPM were widely distributed, involving connections from all macroscale brain regions and canonical networks. The high-attention network was weighted toward prefrontal and motor regions with large contributions of connections involving the frontoparietal, medial frontal, and salience networks. These results fit broadly with accounts of age-related compensatory over-recruitment of bilateral frontal areas (e.g., Park & Reuter-Lorenz, 2009), providing converging evidence, from a network perspective, that increased connectivity among frontal areas supports attentional performance in older adults. Indeed, one previous study illustrated that older adults who demonstrated increased connectivity between bilateral prefrontal areas performed similarly to young-adult comparators, whereas the group that did not exhibit this increase performed much lower (Eavani et al., 2016). The large degree of frontal network involvement in the high-attention network is also supported by a robust literature documenting the role of frontally-mediated networks as “hubs” that regulate other brain networks (Cole et al., 2013; Spreng et al., 2010) in service of memory, attention, and executive function (Grady et al., 2016; La Corte et al., 2016; Shaw et al., 2015). For example, connectivity between the salience network with other networks was predictive of performance on tasks of episodic memory, working memory, and inhibition (La Corte et al., 2016). Moreover, frontoparietal network connectivity mediates the relationship between connectivity in other networks and performance, supporting its role as a primary driver of cognitive function in older adults (Shaw, Schultz, Sperling, & Hedden, 2015).

In contrast, the low-attention network was most heavily weighted towards subcortical areas, with large contribution of subcortical-medial frontal and visual II-medial frontal connections. Interestingly, previous CPM analyses using the gradCPT have also found a similarly large contribution of connections involving subcortical areas to the low-attention network (Rosenberg, Finn, et al., 2016), suggesting that this may be a consistent feature of poor attentional performance. When comparing the high- and low-attention networks, there is a notable lack of inter-frontal network (i.e., frontoparietal-medial frontal) contribution to the low-attention network, again pointing to a supportive role of frontal involvement for attention in older adults. Whereas connectivity within several expected networks (i.e., medial frontal, frontoparietal, and salience networks) predicted better attention, many of the edges of the low-attention network were between disparate networks (e.g., subcortical-medial frontal). This is suggestive of the well-characterized pattern of diffuse increases in between-network connectivity in aging (Ferreira et al., 2016), which can result in the merging of previously modular networks (Geerligs, Renken, et al., 2015). This loss of functional segregation has been found to be detrimental to cognition for older adults (Andrews-Hanna et al., 2007; Avelar-Pereira et al., 2017; Damoiseaux et al., 2008; Geerligs, Renken, et al., 2015; Geerligs et al., 2014; Grady et al., 2016; Onoda et al., 2012), and consistent with our observation of such patterns in a network predicting poorer attentional performance.

Although the Aging-CPM was ultimately generalizable across samples, it is important to consider factors that may have hampered initial model derivation success. Across rounds of cross-validation, we observed considerable variability in the edges selected as relevant to performance, suggesting that there were heterogeneous patterns of brain-behavior relations across the sample. Decades of research have demonstrated that complex changes in brain structure and function result in diverse profiles of cognitive function in later life, with some linked to progressive decline over time while others having little impact (Boyle et al., 2017). Therefore, even samples of non-cognitively impaired older adults likely exhibit varied presence and extent of neuropathology and functional alterations. It may be the case that the sample used for model derivation in this study was not sufficiently sized (internal validation included *N*_1_ = 41) for the model to converge on a solution in the face of such variability. Importantly, other studies investigating CPM models in larger aging datasets have also encountered mixed success, both in attempting model derivation (Lin et al., 2018) and in testing generalizability of previously identified networks (Avery et al., 2019). This suggests that the heterogeneous process of aging poses a unique challenge to brain-based biomarker identification.

Although age-related differences in functional connectivity are amplified during challenging cognitive tasks compared to rest (Dørum et al., 2017), it is possible that either the gradCPT task or the selection of *d’* as the behavior of interest was not optimal for elucidating individual differences in sustained attention in older adults. Recent studies have shown that certain cognitive states appear to amplify individual differences in cognition and some tasks yield better prediction than others (Greene et al., 2018). Given that the development of useful biomarkers hinges greatly on the metric of behavior it is trained upon, future studies would likely benefit from scanning older adults as they perform a suite of tasks that tap multiple complementary domains of attention. As such, future research may confront these challenges by employing multimodal and multivariate approaches (Cole & Franke, 2017; Yoo et al., 2019), integrating data regarding structural features and connectomes from both resting-state and various tasks.

Despite the challenges posed by heterogeneity in this aging sample, the present study produced a model that explained significant variance in attentional performance. This suggests that even in the face of age-associated neuropathology and heterogeneous brain-behavior relationships, there is a pattern of connectivity capturing attentional abilities that is shared, which can successfully predict performance in an independent sample of older adults. These results highlight the potential utility of brain-based models for predicting individual-level attentional functioning in the context of the heterogeneous aging process. Future work utilizing multimodal and multivariate approaches to capture this variability (Yoo et al., 2019) holds promise for producing imaging-based biomarkers that are sensitive predictors of current cognitive status. These biomarkers would offer exciting avenues for predicting future cognitive decline and serving as target outcomes for interventions aimed at promoting cognitive function in older adults.

## Notes

**Funding:** This work was supported by the National Institute on Aging of the National Institutes of Health [Award Number: R01AG054427]. The content is solely the responsibility of the authors and does not necessarily represent the official views of the National Institute on Aging or the National Institutes of Health.

**Conflict of Interest:** None to disclose.

### Competing Interest Statement

The authors have declared no competing interest.

