## Supplementary Materials for "Defining a Connectome-Based Predictive Model of Attentional Control in Aging"

**Additional Motion Controls**

Older adults exhibit greater head motion than young adults on average during both task-based and resting-state scans, raising concerns about how to adequately control for these differences in studies of age-related differences (Cao et al., 2014; Geerligs, Renken, et al., 2015; Geerligs, Rubinov, CAM-CAN, & Henson, 2015; Geerligs, Tsvetanov, & Henson, 2017; Van Dijk, Sabuncu, & Buckner, 2012). Although differences in head motion can introduce confounds in studies of functional connectivity, there is increasing evidence that in-scanner motion reflects neurobiological, or trait-level, characteristics that may also be related to true differences in functional connectivity and cognition (Wylie, Genova, DeLuca, Chiaravalloti, & Sumowski, 2014; Zeng et al., 2014). Therefore, attempts to remove motion artifacts, may obscure real and meaningful effects of age, cognitive ability, and behavior on functional connectivity along with artifactual ones (Geerligs et al., 2017; Wylie et al., 2014).

In light of these issues, we followed the recommendations of Geerligs et al. (2017) set forth specifically for studies examining individual differences in functional connectivity in healthy older adults. Recommendations include regression of motion parameters at the participant level; regression of cerebrospinal fluid (CSF) and white matter (WM) signal, found to reduce the effects of age-differences in motion and vascular health; high-pass filtering and pre-whitening; use of a larger smoothing kernel, found to minimize the effects of age-related differences in the location of functional regions; and Pearson correlation to compute functional connectivity estimates. These steps are described in more detail in the *Image Preprocessing* section. Consistent with a well-established pipeline in our laboratory (Fountain-Zaragoza et al., 2019) and a prior study employing CPM in older adults (Lin et al., 2018), participants were excluded for excessive head motion, which was defined as mean framewise displacement (FD) >

.15 mm and motion > .5 mm in more than 10% of functional volumes (Power, Barnes, Snyder, Schlaggar, & Petersen, 2012). Additionally, we corrected for head motion in several ways through nuisance regression as in the Image Preprocessing section.

Following model derivation, we conducted several follow-up analyses to assess the potential confounding effects of head motion. In addition to motion controls implemented during preprocessing and covarying motion at the edge selection step (see Methods), we confirmed that the derived models were not able to predict head motion (full: *rho* = 0.15, *p* = .35; high: *rho* = 0.13, *p* = .42; low: *rho* = 0.15, *p* = .36), and that strength in these networks was not significantly associated with motion (high: *rho* = 0.13, *p* = .42; low: *rho* = -0.15, *p* = .36). However, we did find that the model's predicted *d'* values remained associated with motion (*rho* = 0.38, *p* = .015). Notably, the model's performance improved when conducting a partial correlation between predicted and observed *d'* controlling for mean FD, with successful prediction from the full model (*rho* = 0.32, *p* = .044, permutation *p* = .03) and the low-attention network (*rho* = 0.41, *p* = .008, permutation *p* = .005), but not the high-attention network (*rho* = 0.19, *p* = .24, permutation *p* = .16). These findings suggest that although we were successful in selecting networks of edges whose connectivity was not related to motion, estimates of predicted performance in this sample are improved when controlling for motion.

**Testing Edge Selection Thresholds**

As the choice of significance threshold for edge selection is inherently arbitrary, and in accordance with troubleshooting suggestions in Shen et al. (2017), we attempted derivation using various significance thresholds, as done in previous studies (cf., Gao, Greene, Constable, & Scheinost, 2019; Greene et al., 2018; Jangraw et al., 2018; Yoo et al., 2018). In light of our concerns that the models were overfitting to noise, and in an effort to retain only edges that are most strongly related to behavior across the training sets rather than any edges reaching a certain level of significance, we also chose to evaluate edge selection based on percentiles. Accordingly, we attempted derivation for models selecting edges associated with behavior at *p* values of .1, .05, .01, .005, and .001, and edge-behavior correlation coefficients above the $90^{th}$, $95^{th}$, $99^{th}$, $99.5^{th}$ and $99.9^{th}$ percentiles. Internal validation was not successful for any of these hyperparameters (Table S1).

Varying threshold values resulted in vastly different network sizes—with the strictest thresholds identifying networks containing only 23 – 75 edges and the most lenient thresholds yielding networks with over 4000 edges. Additionally, within each threshold value, there was a considerable range in the number of edges selected in any one round of cross-validation (Table S1 "Cross-Round # Edges"). In contrast, the number of edges found in every round of cross-validation (Table S1 "Consistent # Edges") was notably smaller. In fact, for the various threshold types, only 14.7 – 40.2% of edges selected in any round of cross-validation were found consistently across rounds. This provided further evidence that the models were fitting to noise, or features not shared by all members of the internal validation sample, raising the question of whether prediction might be successful based on these shared features alone.

Table S1. *Internal Validation Prediction Results at Various Edge Selection Thresholds.*

| Edge Selection Threshold | | Internal Cross-Validation *rho* | *Sig.* | Cross-Round # Edges | Consistent # Edges | Final Model Internal Prediction[†] *rho* |
|---|---|---|---|---|---|---|
| ***p*-value** | .1 | -0.005 | 0.98 | 3504 – 4187 | 1501 | 0.79 |
| | .05 | 0.069 | 0.67 | 1685 – 2247 | 708 | 0.80 |
| | .01 | 0.16 | 0.31 | 295 – 553 | 122 | 0.77 |
| | .005 | 0.10 | 0.53 | 145 – 325 | 58 | 0.74 |
| | .001 | 0.12 | 0.45 | 23 – 75 | 11 | 0.64 |
| **Percentile** | 90 | 0.001 | 0.99 | 3524 | 1417 | 0.79 |
| | 95 | 0.079 | 0.63 | 1762 | 662 | 0.79 |
| | 99 | 0.16 | 0.32 | 352 | 107 | 0.79 |
| | 99.5 | 0.11 | 0.51 | 176 | 53 | 0.75 |
| | 99.9 | 0.093 | 0.56 | 35 | 9 | 0.66 |

[†] The prediction estimates resulting from creating the final models are measures of in-sample fit that can be inflated by over-fitting due to deriving and testing the models in the same dataset. These results should not be interpreted as true estimates of prediction.

**Alternative Derivation Strategies**

Prediction estimates based on leave-one-out cross-validation (LOOCV) are limited in that they are based on only one participant serving as a test subject in each round. This method also does not permit permutation, as the same networks will be selected on each round of cross-validation. To rule out the possibility that our choice of cross-validation strategy contributed to failed internal validation, we conducted iterations of *k*-fold cross-validation as an alternative approach. We used a 10-fold cross-validation in which on each round of cross-validation, the sample is divided into 10 groups, nine of which are used as the training set and one group is left out as the test set. Since we had an indivisible number of internal validation participants (41), we iteratively left one participant out and conducted 100 iterations of 10-fold cross-validation. This resulted in a distribution of 4100 total model estimates (correlation coefficients between predicted and observed scores) for derivation of the high-attention, low-attention, and combined network (Figure S1). Internal validation remained unsuccessful in nearly all iterations. For the combined model, mean $rho = 0.11$ and mean $p = .49$; for the high-attention model, mean $rho = 0.06$ and mean $p = .65$; for the low-attention model, mean $rho = 0.14$ and mean $p = .39$.

Additionally, given that linear regression can be susceptible to overfitting to training data particularly when the sample size is relatively small and features are highly multicollinear, we tested derivation using ridge regression. Ridge regression employs a regularization parameter ($\lambda$) which penalizes the estimates to reduce the impact of influential features. On each round of LOOCV, feature selection was conducted using the same procedures described in the main text and an optimal $\lambda$ hyperparameter was determined through 10-fold cross-validation using the training set (code adapted from https://github.com/YaleMRRC/CPM/tree/master/matlab/misc). Derivation remained unsuccessful using this approach ($rho = 0.17$, $p = .28$).

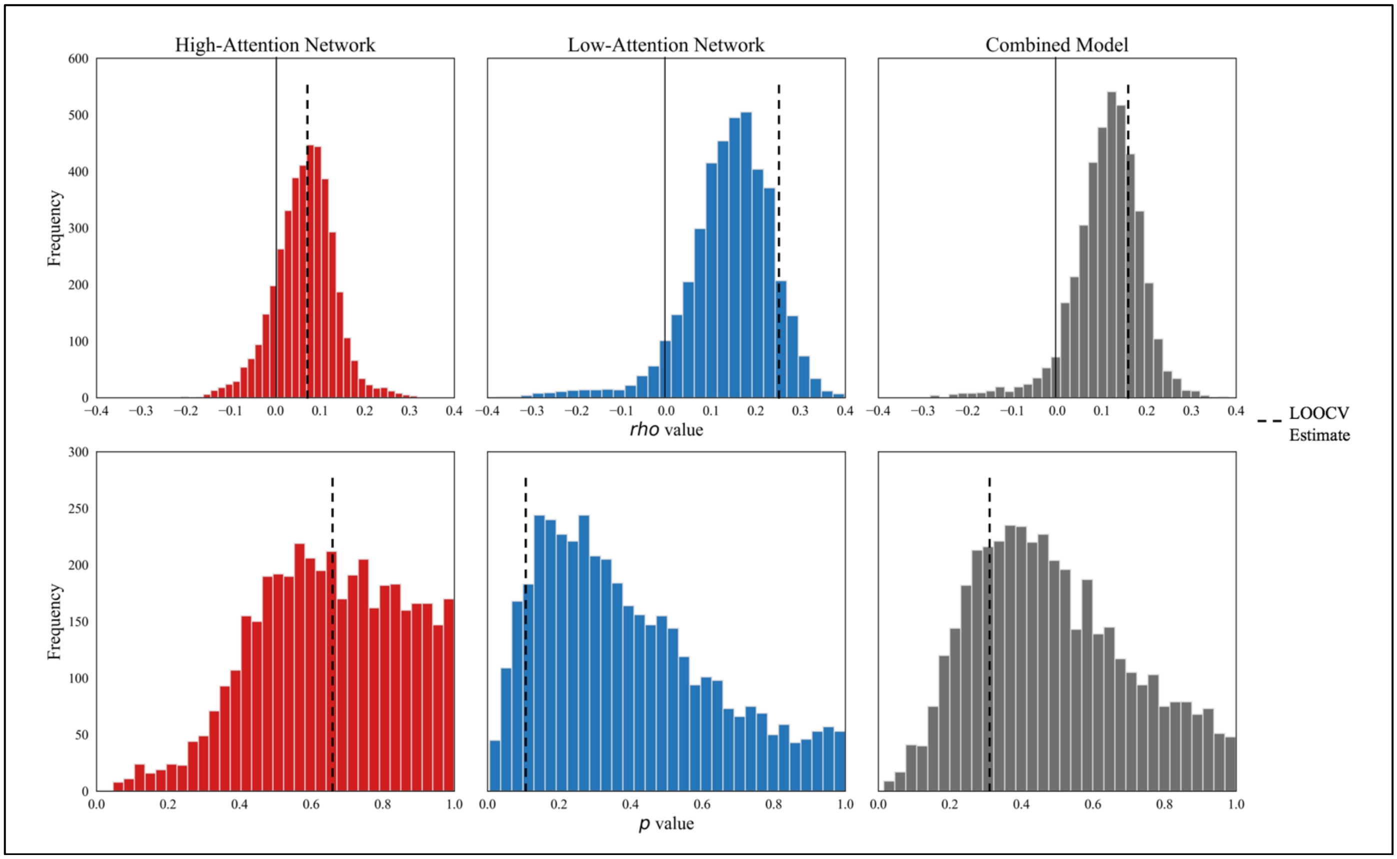


Figure S1. Permuted derivation results. Histograms depict the distribution of *rho* (top row) and *p*-values (bottom row) of 4100 iterations of *k*-fold cross-validation. Results are shown for the high-attention network (red, left column), low-attention network (blue, middle column), and combined model (gray, right column). The vertical dashed line on each graph shows the model estimates from leave-one-out cross-validation (LOOCV; presented in the main text) for comparison.

### External Validation at Multiple Thresholds

The external validation results for each threshold are presented in Figure S4; however, note that models with thresholds of $p < .001$ and 99.9$^{th}$ percentile are not plotted because they yielded a very small number of consistent edges and were markedly less correlated with *d'* in the internal validation sample relative to all other models. Models based on more stringent thresholds accounted for 30.2% ($p = .005$) at $p < .005$, 31.6% ($p = .014$) at 99.5$^{th}$ percentile, 21.2% ($p = .012$) at $p < .001$, and 30.6% ($p = .014$) at 99.9$^{th}$ percentile. Models employing more lenient thresholds were less successful, accounting for 15.5% at $p < .1$ ($p = .028$), 16.7% at 90$^{th}$ percentile ($p = .029$), 15.7% at $p < .05$ ($p = .026$), and 16.1% at 95$^{th}$ percentile ($p = .027$).

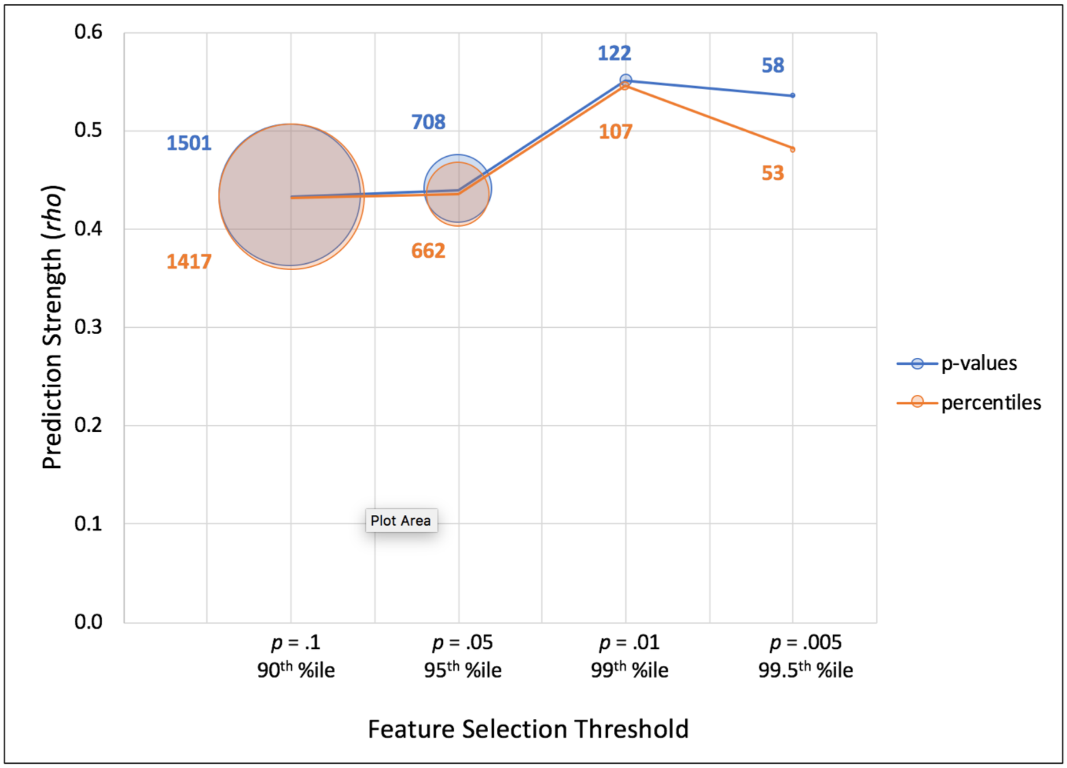


Figure S2. External predictive ability for models at various edge selection thresholds. Models using *p*-value thresholds are presented in green and those using percentile thresholds are presented in orange. Network size is proportionate to the diameter of each circle, and the number of consistent edges found across rounds of cross-validation are denoted next to each circle in the corresponding color. Predictive power was highest for models using thresholds of $p = .01$ and 99$^{th}$ percentile.
